# Disease-modifying effects of Vincamine supplementation in *Drosophila* and human cell models of Parkinson’s disease

**DOI:** 10.1101/2022.12.28.522104

**Authors:** Francisco José Sanz, Cristina Solana-Manrique, Nuria Paricio

## Abstract

Parkinson’s disease (PD) is an incurable neurodegenerative disorder caused by the selective loss of dopaminergic neurons in the *substantia nigra pars compacta*. Current therapies are only symptomatic, and are not able to stop or delay its progression. In order to search new and more effective therapies, our group carried out a high-throughput screening assay, identifying several candidate compounds able to suppress motor defects in *DJ-1β* mutant flies (a *Drosophila* model of familial PD) and to reduce oxidative stress (OS)-induced lethality in *DJ-1*-deficient SH-SY5Y human cells. One of them was vincamine (VIN), a natural alkaloid obtained from the leaves of *Vinca minor*. Our results showed that VIN is able to suppress PD-related phenotypes in both *Drosophila* and human cell PD models. Specifically, VIN reduced OS levels in PD model flies. Besides, VIN diminished OS-induced lethality by decreasing apoptosis, increased mitochondrial viability and reduced OS levels in *DJ-1*-deficient human cells. In addition, we have demonstrated that VIN is able to exert its beneficial role, at least partially, by the inhibition of voltage-gated Na^+^ channels. Therefore, we propose that these channels might be a promising target in the search for new compounds to treat PD, and that VIN constitutes a potential therapeutic treatment for the disease.

## 1. INTRODUCTION

Parkinson’s disease (PD) is a progressive and incurable neurological disorder caused by the selective loss of dopaminergic (DA) neurons in the *substantia nigra pars compacta*, which leads to reduced dopamine levels in the striatum (Vázquez-Vélez et al., 2021). However, alterations in other neurons as well as in other brain regions have also been found (Feraco et al., 2021; Koszła et al., 2021). Since neurodegeneration in PD is the result of the combination of processes occurring inside and/or outside the cells, its etiopathogenesis is not yet completely understood. Despite this, several works suggest that mitochondrial alterations, protein misfolding and aggregation, autophagy defects, inflammation, increased oxidative stress (OS) levels, calcium dyshomeostasis, and metabolic alterations might play an important role in the development of the disease (Anandhan et al., 2017; Groenendyk et al., 2021; Mahmood et al., 2021; Poewe et al., 2017; Solana-Manrique et al., 2022). PD is characterized by a range of motor symptoms such as bradykinesia, resting tremor, rigidity and postural instability. Besides, PD also curses with non-motor symptoms including mood alterations, sleep disturbances or even dementia, which significantly reduce patients’ quality of life (Feraco et al., 2021; Gouda et al., 2022; Mahmood et al., 2021).

Current options to treat PD are limited, and are mainly based on restoration of dopamine levels in the striatum (Majali et al., 2021). These approaches represent the standard treatment for motor symptoms, but they are not able to halt or delay the progression of the disease (Gouda et al., 2022; Majali et al., 2021). PD is the second most common neurodegenerative disease affecting 1-2% of people over the age of 65, a percentage expected to increase in the near future (Gouda et al., 2022; Mahmood et al., 2021). In fact, PD is currently the fastest growing neurological disorder in the world (Fletcher et al., 2021). Therefore, there is an urgent unmet medical need for the identification and development of novel and effective therapies to treat this disease. Several experimental approaches are being used to achieve these goals like gene therapy, immunotherapy, the use of neurotrophic factors, stem cell therapy or the design of high-throughput screening (HTS) platforms and drug repurposing strategies (Aldewachi et al., 2021; Balakrishnan et al., 2021; Fletcher et al., 2021; Sanz et al., 2021; Stoker & Barker, 2020; Tomishima & Kirkeby, 2021), among others. In this scenario, we have recently performed an in vivo HTS assay aimed to identify new potential candidate compounds to treat PD, using a *Drosophila* model of the disease based on inactivation of the *DJ-1β* gene (the fly ortholog of human *DJ-1*, a gene involved in familial PD cases) (Sanz et al., 2021). *DJ-1β* mutant flies exhibit several PD-related phenotypes such as locomotor defects, reduced lifespan as well as increased OS levels (Lavara-Culebras et al., 2010; Lavara-Culebras & Paricio, 2007; Sanz et al., 2023). Among the 1120 drugs included in the Prestwick® chemical library, we identified ten compounds not only able to suppress motor defects of PD model flies but also to reduce OS-induced lethality in *DJ-1*-deficient SH-SY5Y human cells; therefore, these drugs constitute promising therapeutic agents for PD (Sanz et al., 2021). One of the compounds identified was vincamine (VIN) (referred to as compound B in that study), a natural alkaloid obtained from the leaves of *Vinca minor*, commonly known as lesser or dwarf periwinkle (Fayed, 2010). Several studies have indicated the potential role of nutraceuticals, such as VIN as well as its semi-synthetic derivative vinpocetine, to target the underlying neurodegenerative processes of PD (Lama et al., 2020).VIN exhibits antioxidant and anti-inflammatory activities, and may work through several mechanisms of action. It is a phosphodiesterase (PDE) I inhibitor, a blocker of voltage-gated Na^+^ channels (VGNC), and a GPR40 agonist (Abdel-Salam et al., 2016; Du et al., 2019; Sheref et al., 2021). VIN is commercially available in the United States as a health care product with nootropic function, and exerts a beneficial effect in a number of brain disorders of elderly patients, such as memory disturbances, vertigo, transient ischemic deficits and headache (Fandy et al., 2016). It enhances cerebral blood flow and glucose uptake, and it is prescribed to treat memory deficits and cognitive impairments also in Alzheimer’s disease (AD) patients (Abdel-Salam et al., 2016). However, VIN has been barely tested in animal models as candidate compound to treat PD. Only a recent study has shown that VIN administration reduced motor defects and OS levels in a haloperidol-induced rat PD model and exerted an anti-inflammatory effect (Sheref et al., 2021).

In this work, we have evaluated the therapeutic potential of VIN in several PD models. Our results have demonstrated that VIN suppressed PD-relevant phenotypes in *Drosophila* and human cell PD models based on *DJ-1* deficiency such as high OS levels, overactivation of the pre-apoptotic JNK pathway and mitochondrial dysfunction. In addition, we have found that VIN could be exerting a neuroprotective effect through the blockage of VGNC, thus supporting that these channels might be a promising target in the identification of new treatments for PD.

## 2. MATERIALS AND METHODS

### 2.1. Drosophila stocks

*DJ-1β* mutant flies from the *DJ-1β*^*ex54*^ strain (Park et al., 2005) were used in this study. Flies were maintained and cultured in standard *Drosophila* medium at 25 °C, unless otherwise indicated.

### 2.2. Quantification of H_2_O_2_ levels and protein carbonyl group formation in fly extracts

H_2_O_2_ and protein carbonylation levels were measured in 5-day-old *DJ-1β* mutant flies treated with vehicle (0.1% DMSO) or with 10 μM vincamine. Quantification of H_2_O_2_ levels was carried out in fly extracts using the Amplex Red Hydrogen Peroxide/Peroxidase Assay Kit (Invitrogen) as previously described in (Sanz et al., 2017). Protein carbonyl groups were quantified in fly extracts using 2,4-dinitrophenyl hydrazine derivatization as previously described in (Sanz et al., 2021). All experiments were carried out using three biological replicates and three technical replicates for each sample.

### 2.3. SH-SY5Y cells culture and drug treatment

In this study, we used *DJ-1*-deficient and *pLKO*.*1* control SH-SY5Y neuron-like cells previously generated by our laboratory (Sanz et al., 2017). Cells were cultured at 37 °C and 5% CO_2_ in selective growth medium consisting of Dulbecco’s Modified Eagle Medium/Nutrient Mixture F-12 (DMEM/F12) (Biowest) and supplemented with 10% (v/v) fetal bovine serum (Capricorn), 1% non-essential amino acids, 100 mg/ml penicil/streptomycin (Labclinics) and 2 μg/mL puromycin (Labclinics). Viability of cells treated with vincamine, the VNGC activator veratridine, or DMSO 0.1% as vehicle was evaluated using a MTT (3-(4, 5-dimethylthiazol-2-yl)-2-5-diphenyltetrazolium bromide) (Sigma-Aldrich) assay as described in (Sanz et al., 2017). To determine whether veratridine was able to interfere with the neuroprotective effect of our candidate compound, cells were pretreated for 2 h with 150 μM of the VNGC activator before the addition of vincamine. Subsequently, viability assays were performed as described in (Sanz et al., 2017). All experiments were carried out using three biological replicates and three technical replicates for each sample.

### 2.4. Western Blotting

Protein extraction and Western blot of lysates from *DJ-1*-deficient SH-SY5Y cells grown under OS conditions and treated with 10 μM vincamine or with vehicle (0.1% DMSO) were carried out as previously described in (Solana-Manrique et al., 2020). The primary antibodies used in this study were anti-JNK, and anti-phospho-JNK (Thr183/Tyr185) (1:1000, Cell Signaling). The secondary antibody used was anti-rabbit HRP-conjugated (1:5000, Sigma). Quantifications of protein levels were performed with an ImageQuant™ LAS 4000mini Biomolecular Imager (GE Healthcare), and images were analyzed with ImageJ software (NIH). All experiments were carried out using four biological replicates.

### 2.5. Mitochondrial viability

Mitochondrial viability of *pLKO*.*1* control and *DJ-1*-deficient SH-SY5Y cells was evaluated using the MitoTracker™ Red FM (Invitrogen) fluorescence dye as described in (Sanz et al., 2021). Cells were grown in OS conditions (induced with 100 μM H_2_O_2_) and treated either with 10 μM vincamine or with vehicle (0.1% DMSO). Images were acquired using a fluorescence microscope (Leica DMI3000 B), and analyzed with ImageJ software (NIH). All experiments were carried out using three biological replicates and three technical replicates for each sample.

### 2.6. Quantification of O^2-^ levels in human SH-SY5Y cells

Quantification of O^2-^ levels in *pLKO*.*1* control and *DJ-1*-deficient SH-SY5Y cells treated with 10 μM vincamine or with vehicle (0.1% DMSO) was carried out using the dihydroethidium fluorescence dye (Invitrogen), and following a protocol adapted from (Kim et al., 2012). Briefly, 1.8 × 10^4^ cells/well were seeded in a black 96-well plate and incubated for 24 h at 37 °C. Subsequently they were incubated with 100 μM H_2_O_2_ for 3 h. Finally, dihydroethidium was added to each well at a final concentration of 10 μM. Fluorescence was measured at 0 and 30 min at a wavelength of excitation and emission of 540 nm and 595 nm, respectively, in an Infinite 200 PRO reader (Tecan). O^2-^ levels of each sample were calculated using the following formula: [(F30min -F0min) / F0min] × 100. All experiments were carried out using three biological replicates and three technical replicates for each sample.

### 2.7. Statistical analyses

The significance of differences between means was assessed using a t-test when two experimental groups were analyzed. In experiments in which more than two experimental groups were used, the statistical analysis was made using the ANOVA test and Tukey’s post hoc test. Differences were considered significant when P < 0.05. Data are expressed as means ± standard deviation (s.d.).

## 3. RESULTS AND DISCUSSION

### 3.1. Vincamine reduces OS levels in *DJ-1β* mutant flies

Amongst the numerous functions ascribed to the DJ-1 protein, it stands out for its essential role in the defense against OS (Raninga et al., 2017). Increased OS levels are observed in brains of PD patients (Poewe et al., 2017), which suggests that they play an important role in the development of the disease (Poewe et al., 2017; Sanz et al., 2017). According to this, previous studies performed by our group already showed that compounds with antioxidant properties were able to suppress PD-related phenotypes in fly and cell models of the disease based on *DJ-1* deficiency (Casani et al., 2013; Sanz et al., 2017, 2021). As mentioned above, VIN was one of the lead compounds identified in an in vivo HTS assay using *DJ-1β* mutant flies (a *Drosophila* PD model) and validated in *DJ-1*-deficient neuron-like cells (Sanz et al., 2021). Previous studies showed that VIN administration in control rats resulted in a significant reduction of brain iron levels (Fayed, 2010). Iron appears to accumulate in high concentration in neurodegenerative diseases (ND), such as PD or AD, thus contributing to OS and in turn leading to neurodegeneration (Fiedler et al., 2007). Since age-linked ND are characterized by a disturbance in trace element levels in the brain, it was suggested that VIN might exert a beneficial effect in aged people by decreasing OS (Fayed, 2010). It was also shown that VIN was able to reduce Aβ-induced cytotoxicity in PC12 cells by decreasing the concentrations/activities of a variety of OS indicators (Han et al., 2017). As previously reported, *DJ-1β* mutants exhibited high levels of reactive oxygen species (ROS) and protein carbonylation (a post-translational modification caused by high ROS levels) when compared to control flies (Casani et al., 2013; Lavara-Culebras et al., 2010). Indeed, we demonstrated that they had a causative role in motor deficits exhibited by PD model flies (Sanz et al., 2017). In such a scenario, we decided to evaluate whether VIN supplementation could decrease the levels of OS indicators in PD model flies. As shown in Figure 1, we found that *DJ-1β* mutant flies treated with 10 μM VIN during development and 5 days after eclosion presented a significant reduction of H_2_O_2_ (a component of the total ROS pool), and of protein carbonylation levels compared to flies treated with vehicle (Fig. 1). Therefore, our results support that VIN treatment exerts an antioxidant effect in PD model flies based on *DJ-1β* deficiency. Interestingly, a recent study has demonstrated that VIN administration also reduced motor defects and OS levels in a haloperidol-induced rat PD model (Sheref et al., 2021). Taken together, these results support the therapeutic potential of VIN in animal models of PD.

**Figure 1.**
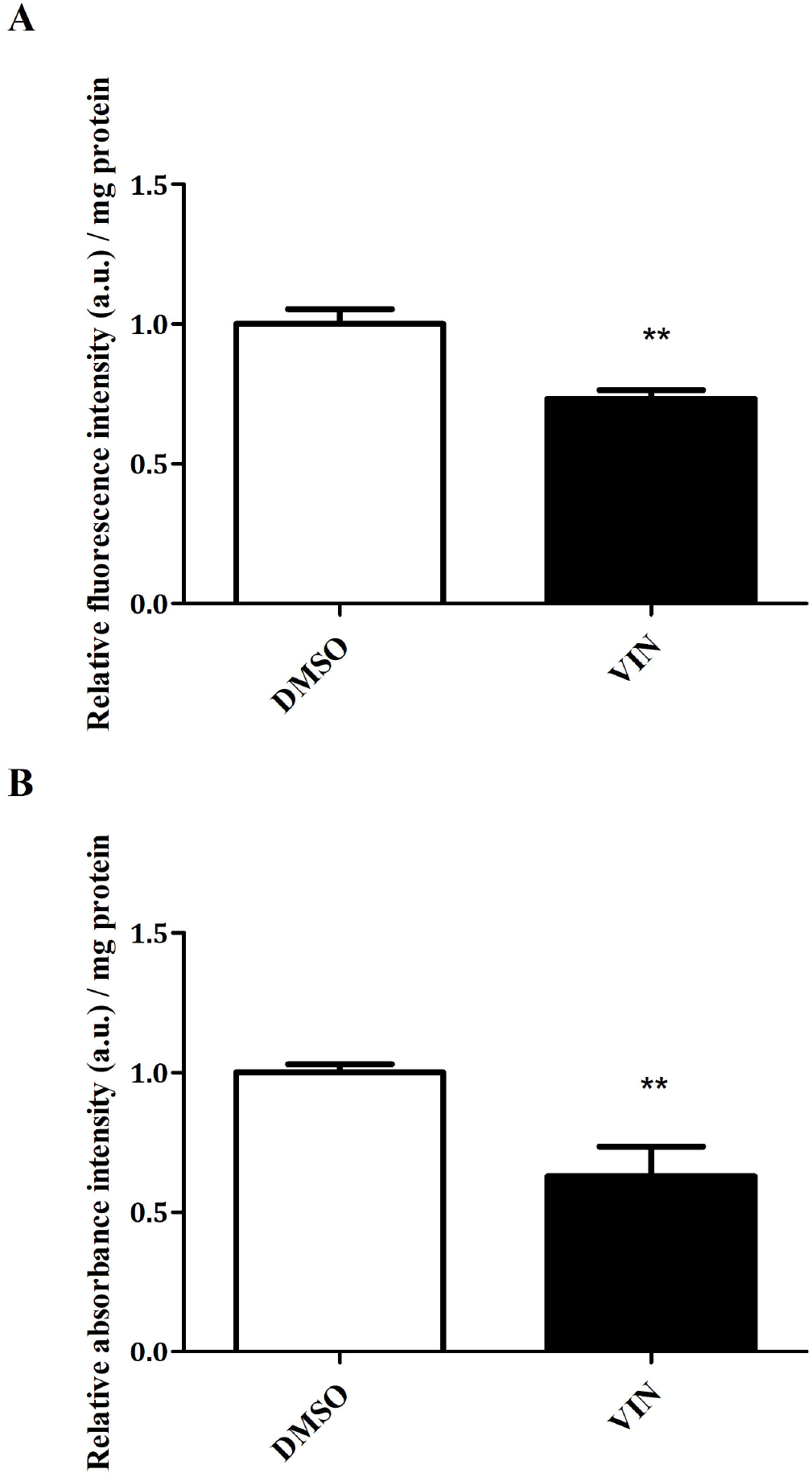
Effect of vincamine on oxidative stress marker levels in *DJ-1β* mutant flies. A) H_2_O_2_ levels in *DJ-1β* mutant flies treated with 10 μM vincamine (VIN) were analyzed using the Amplex H_2_O_2_ Red Kit (Invitrogen). B) Protein carbonylation levels in *DJ-1β* mutants treated with 10 μM vincamine (VIN) were analyzed by absorbance. In all cases, data were expressed as arbitrary units (a.u.) per mg of protein, and results were referred to data obtained in flies cultured in vehicle medium (DMSO). Error bars show s.d. from at least three replicates and three independent experiments (**P < 0.01).

### 3.2. Vincamine increases viability of *DJ-1*-deficient human cells by reducing JNK signaling activation

Although *Drosophila* is an outstanding model organism in the search of new treatments for human diseases, candidate compounds identified in flies have to be validated in mammalian models (Millet-Boureima et al., 2021; Sanz et al., 2021). Previous studies showed that viability of *DJ-1*-deficient human neuroblastoma cells was reduced when cultured in OS conditions (Sanz et al., 2017). We already demonstrated that pretreatment with 10 μM VIN significantly attenuated OS-induced death in *DJ-1*-deficient cells (Sanz et al., 2021). Therefore, we decided to analyze further the effect of VIN in cell viability by pretreating such cells with different concentrations of the compound from 0.1 to 80 μM. Our results showed that VIN exerted neuroprotective effects in a range of 2.5-20 μM, being 10 μM the most effective concentration (Fig. 2A). Therefore, we used this concentration in subsequent experiments performed in human cells. As shown in Fig. 2B, VIN did not affect viability in *pLKO*.*1* control cells at the same concentrations (Fig. 2B).

**Figure 2.**
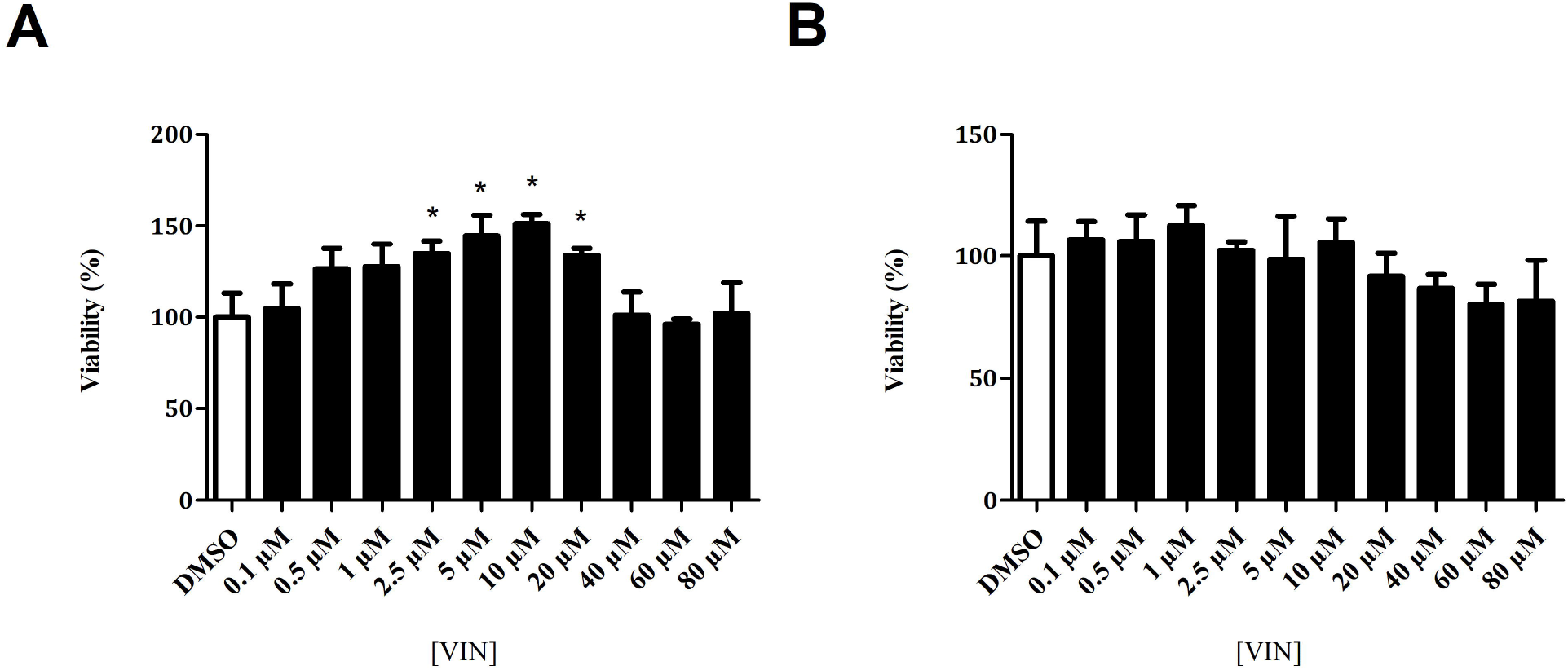
Effect of vincamine on viability of *DJ-1*-deficient and control SH-SY5Y cells. MTT assays were performed to measure the viability of A) *DJ-1*-deficient and B) *pLKO*.*1* control cells grown under OS conditions (induced with 100 μM H_2_O_2_) either treated with vehicle (DMSO) or with different concentrations of vincamine (VIN) (0.1–80 μM). Results were normalized to data obtained in vehicle-treated cells. Error bars show s.d. from three independent experiments in which three biological replicates were used (*P < 0.05).

PD is caused by the loss of DA neurons; however, the reason of this neurodegeneration is still unknown (Ball et al., 2019). Among the processes that might lead to neuronal death, we find apoptosis (Erekat, 2018; Liu et al., 2019), a highly regulated process where the JNK protein plays a key role (Liu et al., 2019). It has been reported that *DJ-1*-deficient SH-SY5Y cells present high levels of JNK phosphorylation, which activates the JNK signaling pathway and promotes cell death (Liu et al., 2019; Sanz et al., 2021; Zhang et al., 2019). In order to evaluate if VIN could be exerting its neuroprotective effect through reducing apoptosis, we performed a western blot assay to study its effect on JNK phosphorylation. We found that JNK phosphorylation levels were significantly reduced in *DJ-1*-deficient cells pretreated with 10 μM VIN compared to cells pretreated with vehicle (DMSO) (Fig. 3, Fig. S1), therefore reducing the activity of the proapoptotic JNK pathway as well as increasing viability of *DJ-1-* deficient cells. In agreement with our results, it was demonstrated that VIN exerted a protective effect in rat livers treated with tamoxifen, a compound that induces cell death (El-Dessouki et al., 2018). Indeed, treatment of rats with VIN ameliorated tamoxifen induced hepatic cell injury *via* suppressing OS and reducing JNK phosphorylation (El-Dessouki et al., 2018).

**Figure 3.**
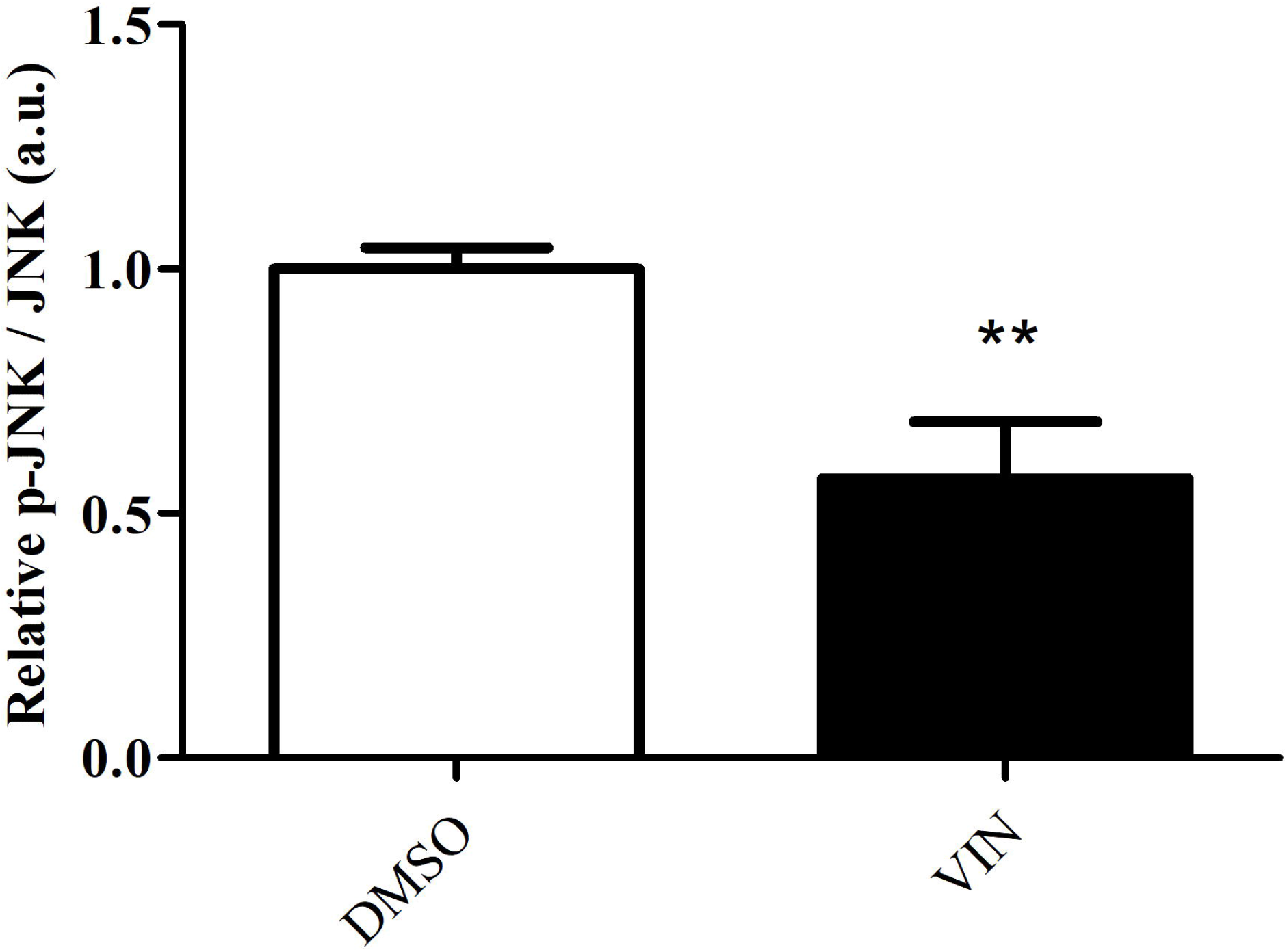
Effect of vincamine on JNK pathway activity in *DJ-1*-deficient SH-SY5Y cells. Western blot analyses were carried out using antibodies against JNK, and p-JNK in *DJ-1*-deficient cells subjected to OS (induced with 100 μM H_2_O_2_) and treated with 10 μM vincamine (VIN). The relative ratio of p-JNK/JNK was analyzed by densitometry. Results were referred to data obtained in vehicle-treated *DJ-1*-deficient cells and expressed as arbitrary units (a.u.). Error bars show s.d. from three independent experiments in which three biological replicates were used (**P < 0.01).

### 3.3. Vincamine enhances mitochondrial viability in *DJ-1*-deficient human cells

Mitochondrial dysfunction plays an important role in PD (Borsche et al., 2021). In fact, many of the genes involved in familial PD cases are functionally associated with mitochondria; this highlights its relevance in PD development (Aryal & Lee, 2019). Specifically, loss of *DJ-1* function has been linked to a reduction of mitochondrial mass as well as to alterations in the morphology and function of this organelle (Chen et al., 2019; Krebiehl et al., 2010; Zhang et al., 2019). Interestingly, the activation of JNK signaling was also related to the onset of mitochondrial dysfunction (Heslop et al., 2020). Previous studies carried out by our group demonstrated that *DJ-1*-deficient human cells presented a reduction of mitochondrial viability compared to *pLKO*.*1* control cells (Sanz et al., 2021). Therefore, we decided to evaluate whether VIN supplementation could increase such viability in mutant cells using the MitoTracker™ Red FM dye. As expected, we found that *DJ-1*-deficient cells presented a significant reduction in the active mitochondrial mass compared to *pLKO*.*1* control cells (Fig. 4A, B). Our results also showed that VIN pretreatment of *DJ-1*-deficient cells resulted in a significant increase of mitochondrial viability compared to those treated with vehicle (Fig. 4A, B).

**Figure 4.**
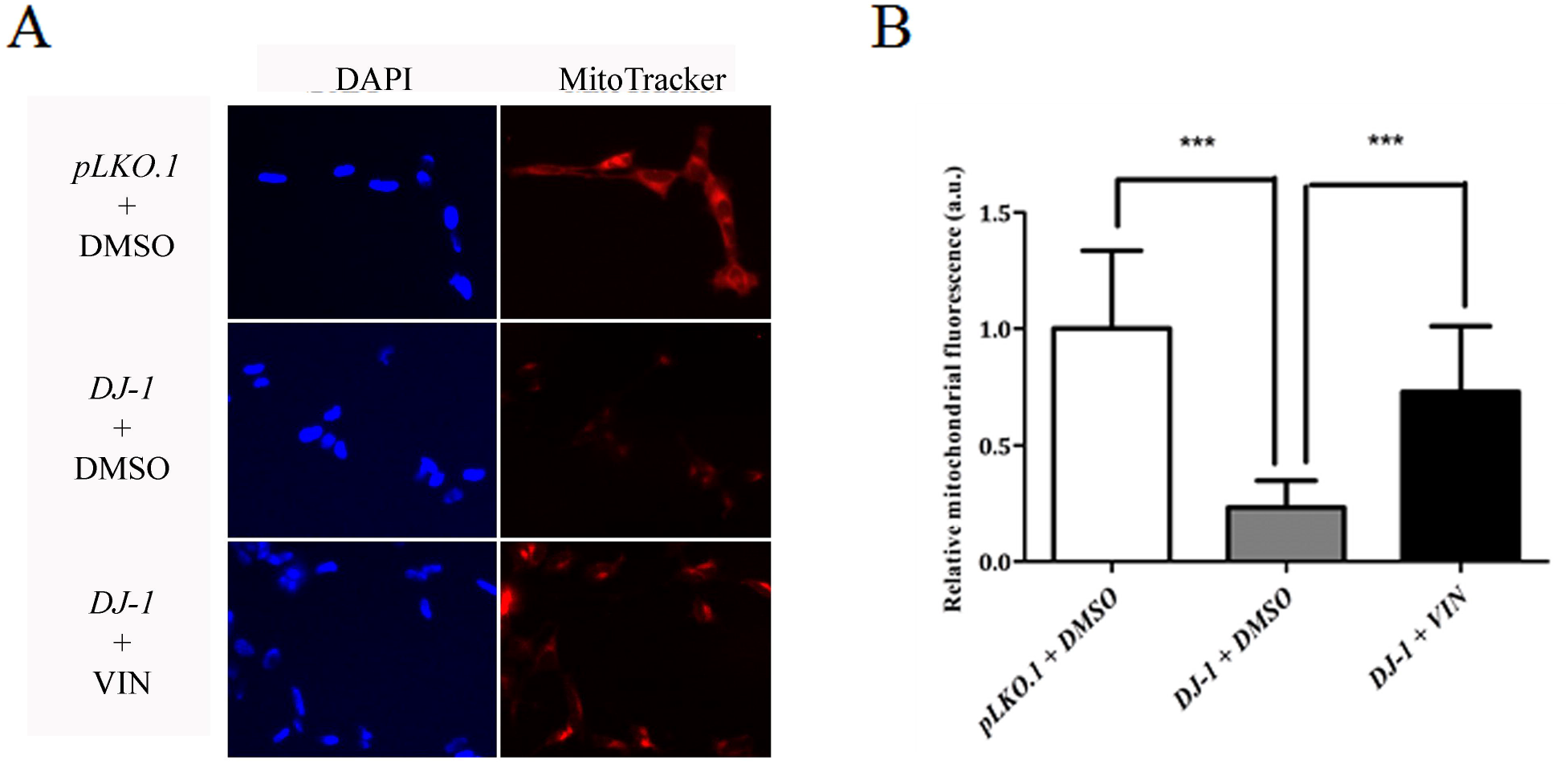
Effect of vincamine on mitochondrial activity in SH-SY5Y cells. A) Representative images of SH-SY5Y cells stained with MitoTracker Red FM, a specific mitochondrial dye, and the nuclear dye DAPI (blue) acquired by fluorescence microscopy. Cells stained were *pLKO*.*1* control cells pretreated with vehicle (DMSO), and *DJ-1-*deficient cells either pretreated with or with 10 μM vincamine (VIN). Scale bar, 50 μm. B) Graphical representation of Mitotracker Red FM fluorescence quantification from A). At least ten images of each strain and condition were analyzed. Results were normalized to data obtained in vehicle-treated *pLKO*.*1* control cells and expressed as arbitrary units (a.u.). Error bars show s.d. from three independent experiments in which three biological replicates were used (***P < 0.001).

Mitochondria are one of the main sources of ROS, which are byproducts of their normal metabolism and homeostasis. Thus, alterations in mitochondrial function may lead to an increase of ROS levels above a toxicity threshold resulting in potentially unwanted oxidative consequences and even cell death (Zorov et al., 2014). For instance, it has been found that mitochondrial damage in PD might be caused by complex I of the electron transport chain (ETC) dysfunction (Aryal & Lee, 2019; González-Rodríguez et al., 2021), a complex to which the DJ-1 protein directly binds (Buneeva & Medvedev, 2021). In addition, several toxins (rotenone, paraquat or MPTP) able to inhibit its activity are commonly used to generate animal and cell models of idiopathic PD (Aryal & Lee, 2019; Wen et al., 2020). Since *DJ-1*-deficient cells showed a decrease in the active mitochondrial mass, we aimed to study the consequence of that reduction in ROS levels. For doing so, we used the dihydroethidium fluorescence dye to quantify O^2-^ levels (a component of the total ROS pool) in *DJ-1* mutant and control cells. We found that *DJ-1*-deficient cells presented higher O^2-^ levels than *pLKO*.*1* control cells (Fig. 5). In addition, we confirmed that *DJ-1*-deficient cells supplemented with VIN showed a reduction in O^2-^ levels compared to vehicle-treated cells (Fig. 5).

**Figure 5.**
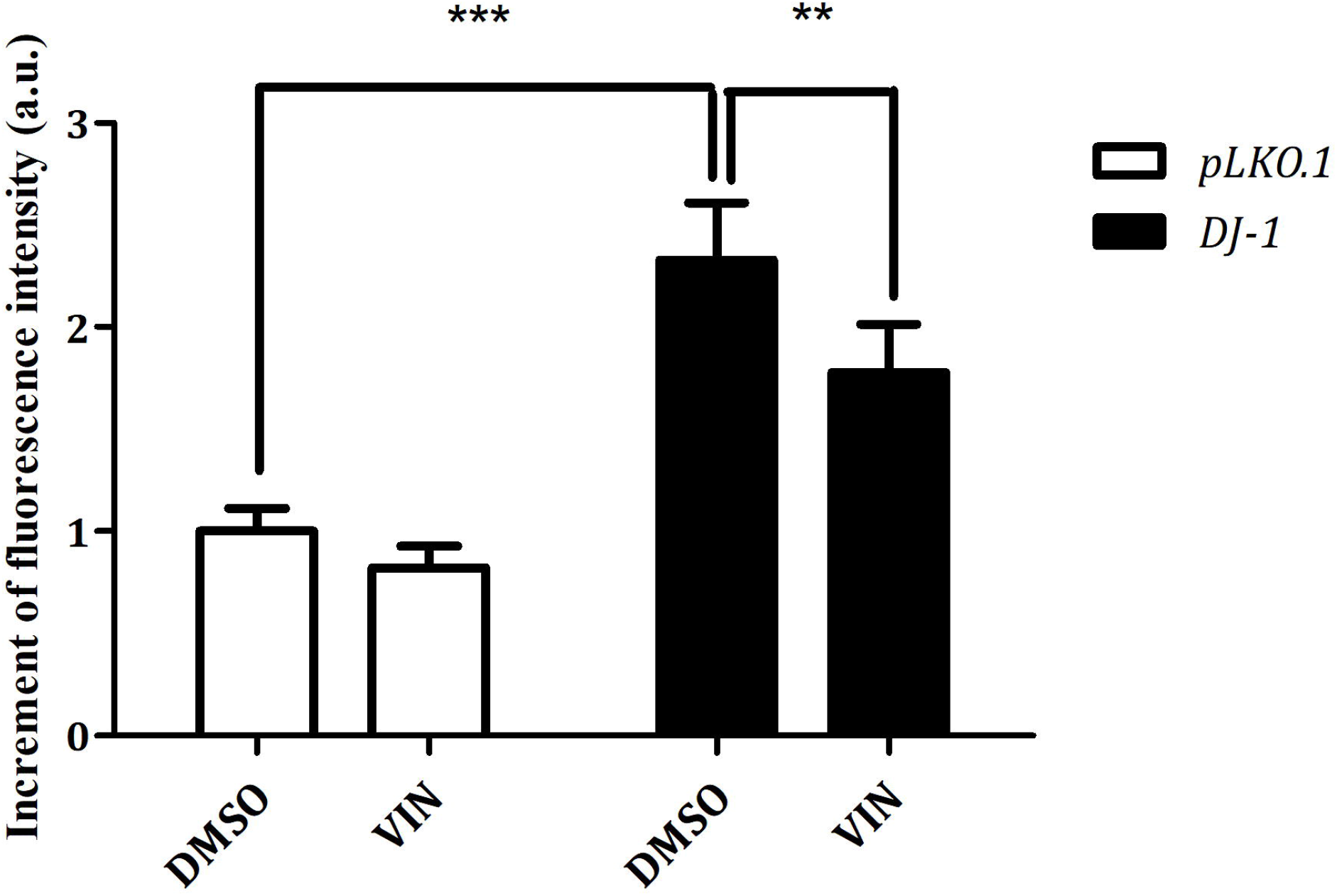
Effect of vincamine on O^2-^ levels in *DJ-1*-deficient SH-SY5Y cells. O^2-^ levels in *pLKO*.*1* and *DJ-1*-silenced cells treated either with vehicle (DMSO) or with 10 μM vincamine (VIN) were analyzed using the fluorescence dye dihydroethidium (Invitrogen). Results were normalized to data obtained in vehicle-treated *pLKO*.*1* control cells and expressed as arbitrary units (a.u.). Error bars show s.d. from three independent experiments in which three biological replicates were used (**P < 0.01; ***P < 0.001).

Taken together, our results support the therapeutic potential of VIN in *DJ*-deficient cells being able to ameliorate PD-associated mitochondrial dysfunction and the consequent increase in ROS levels, as shown in other disease models (El-Dessouki et al., 2018; Sheref et al., 2021).

### 3.4. Veratridine, a VGNCs activator, reduces the neuroprotective effect of vincamine in *DJ-1*-deficient human cells

As previously mentioned, VIN is a compound with multiple mechanism of actions. Several studies have shown that it is a PDE1 inhibitor, a GPR40 agonist, and a blocker of VGNC (Abdel-Salam et al., 2016; Du et al., 2019; Sheref et al., 2021). The effect of PDE inhibitors has been widely studied in several PD models. For example, it was demonstrated that PDE1 inhibitors induced the expression of genes related to neuronal plasticity, neurotrophic factors as well as molecules with neuroprotective function (Medina, 2011). Indeed, PDE inhibitors have been proposed as promising PD therapeutic compounds (Nthenge-Ngumbau & Mohanakumar, 2018). In such scenario, we decided to evaluate whether VIN could be exerting a neuroprotective effect in PD models based on *DJ-1* deficiency through VGNCs inhibition. These channels play a vital role in excitable cells (like cardiomyocytes and neurons) to generate and propagate action potentials. Their functional deficits lead to epilepsy, a brain disorder characterized by seizures and convulsions (Abdelsayed & Sokolov, 2013). Interestingly, a recent study has shown that VGNCs may play an important role in the genesis of cognitive deficits in PD model rats (Wang et al., 2019).

In such a scenario, we aimed to determine if VIN could be exerting a neuroprotective effect in *DJ-1*-deficient cells through inhibition of VGNCs. For doing so, we tested whether veratridine, an alkaloid able to induce persistent activation of these channels (Felix et al., 2004), could affect VIN-mediated neuroprotection. First, we tested different concentrations of veratridine (10-150 μM) in order to identify the maximum concentration of the compound that did not exert a detrimental effect in cell survival. Our results showed that viability of *pLKO*.*1* control and *DJ-1* mutant cells was significantly reduced when using 150 μM veratridine (Fig. 6A, B); in contrast, viability was not affected with lower concentrations of the compound. Therefore, we decided to use 100 μM veratridine to test its effect on cells treated with VIN. Our results showed that viability of VIN-treated *DJ-1*-deficient cells was significantly reduced after veratridine pretreatment (Fig. 6C). Thus, these results indicate that VIN might be exerting a neuroprotective effect, at least partially, through the inhibition of VGNCs. Supporting our results, it was recently reported that VGNCs were overexpressed in a 6-OHDA-induced rat model of PD, and that phenytoin, a VGNC blocker, improved motor and cognitive abilities in that model (Liu et al., 2021). In addition, it was found that treatment with RS100642, a VGNC blocker, reduced levels of OS markers in a rat model of breast cancer (Batcioglu et al., 2012); therefore, VGNC inhibition could play a role in the defense against OS. Overall, these results suggest that VGNCs represents a potential target in the search of novel and more efficient treatments for PD.

**Figure 6.**
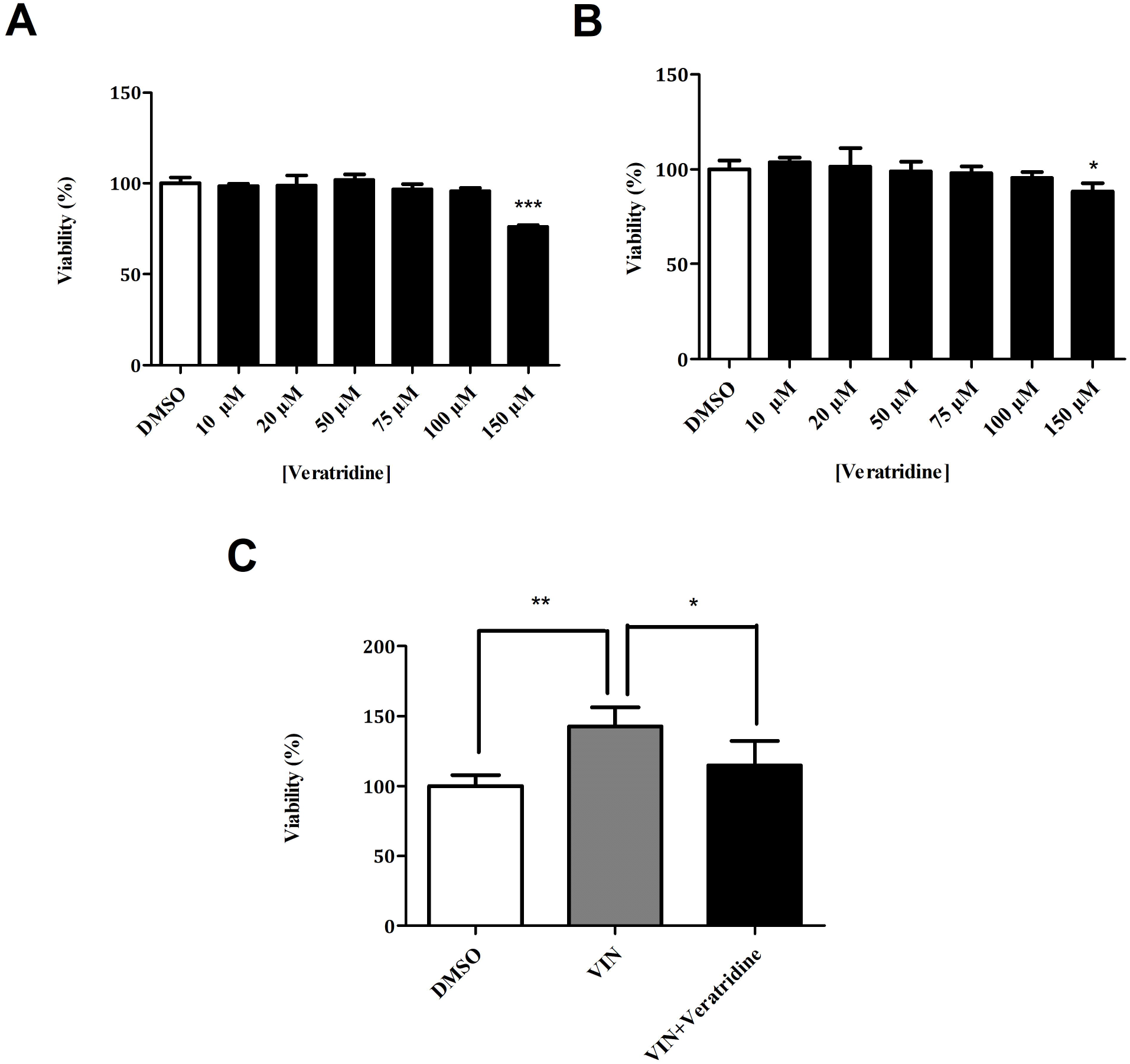
Effect of veratridine on viability of *DJ-1*-deficient SH-SY5Y cells treated with vincamine. MTT assays were performed to measure the viability of A) *pLKO*.*1* control and B) *DJ-1*-deficient cells in the presence of OS (induced with 100 μM H_2_O_2_) and either treated with vehicle (DMSO) or with different with different concentrations of veratridine (1 – 150 μM). In both cases, results were normalized to data obtained in vehicle-treated cells. C) Viability of *DJ-1*-deficient cells under OS conditions and treated with vehicle (DMSO), 10 μM vincamine (VIN) or 10 μM VIN plus 100 μM veratridine. Results were normalized to data obtained in vehicle-treated *DJ-1*-deficient cells. In all cases, error bars show s.d. from three independent biological replicates (*P < 0.05; **P < 0.01; ***P < 0.001).

## 5. CONCLUSIONS

In this work, we have evaluated the therapeutic potential of VIN, a natural alkaloid, as PD treatment using preclinical models of the disease. This study has helped to understand the molecular mechanism of the drug’s action and to determine the conditions in which VIN can mostly exert its beneficial effect in PD models. Indeed, we have demonstrated that VIN is able to ameliorate PD-related phenotypes in *Drosophila* and human cell models based on *DJ-1* inactivation. Specifically, VIN was able to increase viability in *DJ-1-*deficient SH-SY5Y human cells by reducing apoptosis, to increase mitochondrial viability, and to reduce the level of OS indicators. Taken together, these results clearly show the disease-modifying effect of VIN, and allow us to consider this compound as a promising PD therapy. In addition, we have demonstrated that VIN exerts, in part, its neuroprotective effect through VGNCs inhibition, thus supporting these channels as potential targets in the discovery of new and more effective therapeutics to treat PD.

Although studies with VIN in PD models are scarce, the effect of vinpocetine (a VIN derivative with the same mechanisms of action) has been thoroughly tested in animal models of PD and other human diseases (Zhang et al. 2018). For example, this compound was shown to increase cerebral blood flow, hence improving O_2_ and glucose uptake by neurons, which leads to an increase of ATP levels (Jeon et al., 2010). Besides, it has anti-inflammatory effects in PD patients (Ping et al., 2019), and it was shown to reduce motor defects, cognitive alterations, OS levels and DA neurodegeneration as well as to increase dopamine levels in mouse and rat PD models (Ishola et al., 2018; Zaitone et al., 2012). Our results clearly support the continued investigation of VIN as a potential PD treatment. VIN can cross the blood-brain-barrier, is non-toxic, and is used as a dietary supplement, which makes it a good candidate for clinical trials in PD patients. Moreover, it has multiple beneficial properties not only for neural disorders but also as an anticancer agent (Al-Rashed et al., 2021). All these properties make VIN a promising drug for future exploration to identify the precise mechanisms underlying its beneficial effect in PD.

## Abbreviations

AD: Alzheimer’s disease
DA: dopaminergic
ETC: electron transport chain
HTS: high-throughput screening
ND: neurodegenerative disease
OS: oxidative stress
PD: Parkinson’s disease
PDE: phosphodiesterase
ROS: reactive oxygen species
VGNC: voltage gated Na^+^ channels
VIN: vincamine

## ACKOWLEDGEMENTS

The authors received no financial support for this research.

## SUPPLEMENTARY FIGURE LEGEND

**Figure S1**. JNK and P-JNK expression in *DJ-1*-deficient SH-SY5Y cells treated with vincamine. Representative western blot using anti-JNK and anti-P-JNK antibodies in protein extracts from *DJ-1*-deficient cells either treated with vehicle (DMSO) or with 10 μM vincamine (VIN).

