## Supplementary figures and images for "Disease-modifying effects of Vincamine supplementation in *Drosophila* and human cell models of Parkinson’s disease"

### Supplemental Figure 1

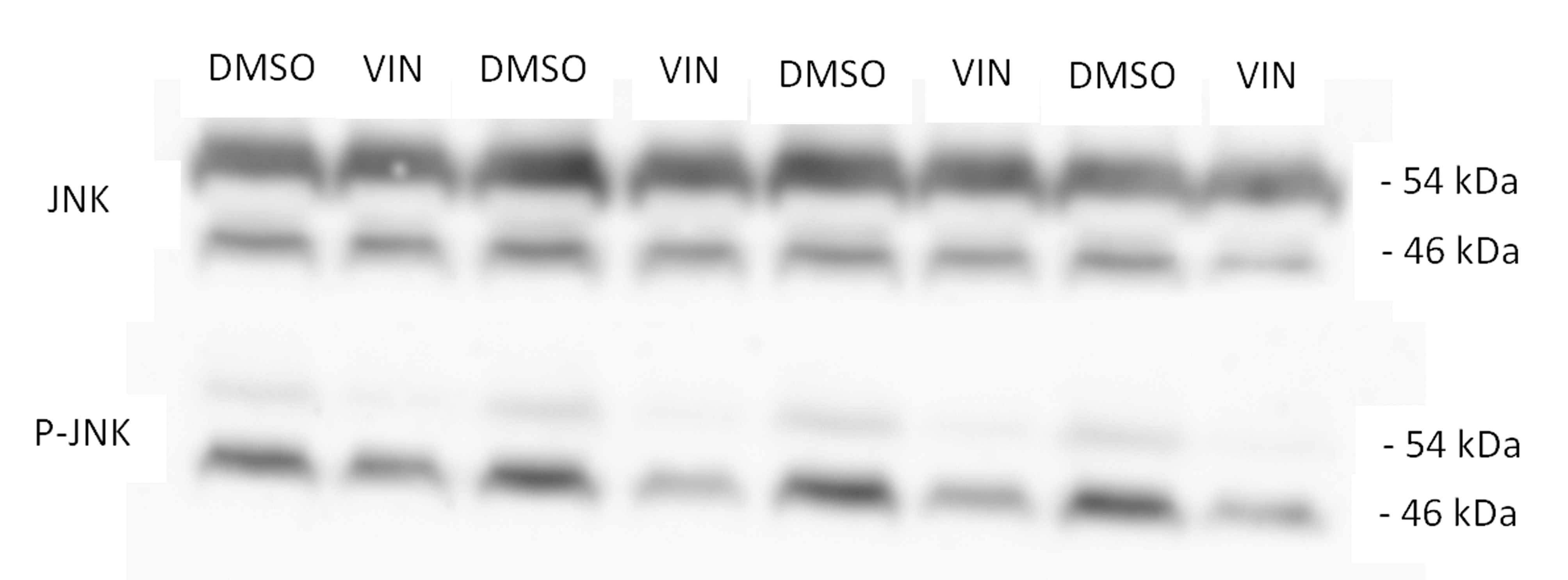
